# Reverse engineering of an aspirin-responsive regulator in bacteria

**DOI:** 10.1101/400788

**Authors:** Lummy Maria Oliveira Monteiro, Letícia Magalhães Arruda, Ananda Sanches Medeiros, Leonardo Martins-Santana, Luana de Fátima Alves, María-Eugenia Guazzaroni, Víctor de Lorenzo, Rafael Silva-Rocha

**Author notes:** Correspondence to: Rafael Silva-Rocha FMRP - Universidade de São Paulo Av. Bandeirantes, 3.900. CEP: 14049-900. Ribeirão Preto, São Paulo, Brazil.

## Abstract

Bacterial transcriptional factors (TFs) and their target promoters are key devices for engineering of complex circuits in many biotechnological applications. Yet, there is a dearth of well characterized inducer-responsive TFs that could be used in the context of an animal or human host. In this work we have deciphered the inducer recognition mechanism of two AraC/XylS regulators from *Pseudomonas putida* (BenR and XylS) for creating a novel expression system responsive to acetyl salicylate (i.e. Aspirin). Using protein homology modeling and molecular docking with the cognate inducer benzoate and a suite of chemical analogues, we identified the conserved binding pocket of these two proteins. Using site directed mutagenesis, we identified a single amino acid position required for efficient inducer recognition and transcriptional activation. While modification of this position in BenR abolishes protein activity, its modification in XylS increases the response to several aromatic compounds, including acetyl salicylic acid to levels close to those achieved by the canonical inducer. Moreover, by constructing chimeric proteins with swapped N-terminal domains, we created novel regulators with mixed promoter and inducer recognition profiles. As a result, a collection of engineered TFs was generated with enhanced response to a well characterized and largely innocuous molecule with a potential for eliciting heterologous expression of bacterial genes in animal carriers.

## Introduction

Transcriptional regulation plays a central role in the adaptation of cells to changing in environmental conditions. In bacteria, this step is mainly regulated by the interaction of RNA Polymerase (RNAP) with the promoter region thought the use of many sigma factors, and by a large number of transcriptional factors (TFs) that can help or block RNAP binding^1^. With the growing interest into the engineering of living cells for novel biotechnological and biomedical application, a special focus has emerged in understanding gene regulation at the molecular level ^1, 2, 3, 4^. In this context, many different classes of TFs have been extensively characterized at the molecular detail from bacteria to mammals and this knowledge has allowed a number of engineering projects, where natural systems can be repurposed to display novel behaviors ^1, 5, 6, 7, 8, 9^. Attempts in this direction are very diverse and examples are the construction of mutated variant of natural TFs with enhanced or modified performance ^10, 11, 12, 13^, the recombination of protein domains to create TFs with completely altered specificity or dynamical behavior ^14, 15^ or the mining of novel regulators from genomes or metagenomes ^16, 17^. Additionally, the revolution provided by CRISPR/Cas9 system has also impacted the field of gene regulation, as this system has been repurposed to construct fully synthetic expression systems based on RNA/DNA interaction ^18, 19, 20^.

Despite the progress on the engineering of novel expression systems, a critical bottleneck relies on the selection of suitable signal-recognition modules related to the application of interest. In other words, while many different TFs are well-characterized as responsive to small molecules (sugars, ions, aromatics, etc.), many times the application at stake requires systems responsive to unusual compounds ^21^. Therefore, construction of TFs variants with enhanced responsivenes to non-natural ligands has become more appealing. Approaches to accomplish this task range from the use of laborious random mutagenesis followed by selection ^11, 12^ to the use of computational analysis to guide rational design ^22^. Here, we focused on the engineering of novel expression systems responsive to commercially available drugs suitable to *in vivo* administration to a mammalian animal. Our particular interest was focused on acetylsalicylic acid (ASA or aspirin), a longstanding, safe and widely used drug. This compound has been applied in synthetic regulatory circuits to deliver lytic proteins to tumors in vivo, representing a promising field for the develo*Pm*ent of tumor-targeting circuits for clinical applications ^23^. As a starting point, we sought to investigate the molecular mechanisms of signal recognition by two homologous regulators of *Pseudomonas putida*, namely BenR and XylS. These two TFs are members of the AraC/XylS family of transcriptional activators ^24, 25, 26^ that recognize different aromatic compounds with structural similarities to ASA. Yet, while both regulators share around 60% amino acid (aa) identity, BenR is responsive to benzoate-only ^25^ while XylS can recognize a large number of substituted variants ^27^. Additionally, some crosstalk between these two regulators and their target promoters has been characterized, as XylS can only recognize its target *Pm* ^24^ while BenR can efficiently activate its natural target *Pb* as well as *Pm* ^25^.

In this we have investigated the molecular mechanisms of signal recognition by these two regulators using computational tools and *in vivo* validation. By constructing a model for the ligand-binding domain of BenR and performing molecular docking with benzoate and a collection of analogues, we identified a potential binding pocket strongly conserved between these two TFs. Thereby, we were able to validate the pocket using site directed mutagenesis and also by constructing chimeric proteins. Finally, we demonstrated how a single aa position plays a critical and opposite role in the activity of both proteins. Some changes of this position in BenR resulted in complete loss of protein activity while the same in XylS triggered an enhanced response to benzoate analogues, including ASA. The results below thus provide not only insights on the mechanism of signal recognition by members of the AraC/XylS family but also the engineering of a regulatory device responsive to aspirin.

## Results

### Analysis of conserved elements in BenR and XylS close homologues and target promoters

In order to investigate the molecular mechanisms accounting for the functional differences between BenR and XylS, we analyzed the close homologues of these proteins existing the genome of some species of *Pseudomonas*. As represented schematically in **Fig. 1**, these proteins are TFs formed by two domains, one N-terminal (AraC domain) which is required for ligand recognition ^28^ and one C-terminal domain composed by two helix turn helix (HTH) regions required for the recognition of the distal and proximal operators (*Od* and *Op*) upstream of the target promoters ^25, 29^. It is proposed that two monomers of XylS are required for the activation of the *Pm* promoter, each binding to an operator region and contacting an A and B box conserved within this region. A previous study ^30^ used alanine scanning mutagenesis to identify four residues in XylS required for the recognition of the A boxes (Arg242, Asn246, Glu249 and Lys250) and five required for the interaction with boxes B (G295, Arg296, Asp299, Asn300 and Arg302). As can be observed in the protein alignment between BenR and XylS homologues, most of these positions are well conserved in the proteins analyzed, with an exception for this been Asn246, Lys250 and Asn300 (**Fig. 1A**). It is surprising to notice that, while the change of Asn to Ala at position 246 in XylS reduced by a half the capability of this protein to activate *Pm*, an Ala is found well-conserved in most BenR protein analyzed. This indicates that BenR homologues might be less stringent in the interaction at the A box of the target promoter ^25^. In the same direction of this hypothesis, analysis of *Pm* and *Pb* promoters from several *Pseudomonas* strains reveals that most features (A and B boxes, -35/10 region) are well conserved, except for the A box of *Pb* (the target of the BenR studied here) from *Pseudomonas putida* KT2440 (**Fig. 1B**). In this sense, to check the effect of the A box on the interaction between XylS and *Pm* and *Pb* promoters, we assayed the promoter activity using a lux reporter system. We used a wild type xylS gene expressed from a pSEVA vector ^31^ and *Pm, Pb* and *Pb*-syn1 (a *Pb* variant with the recomposed A box of Od region ^25^). Using this system, we observed that while XylS could recognize *Pm* very efficiently, it was not able to induce *Pb* activity in response to the inducers tested (**Fig. 2**). However, when a version of *Pb* with the reconstituted A box (*Pb*-syn1) was used, we could observe a strong induction of promoter activity in response to the inducers used. Taken together, these results reinforce the notion that while XylS has a critical requirement for complete A and B boxes at the Od and Op regions, BenR is less stringent for promoter recognition ^25^.

**Figure 1.**
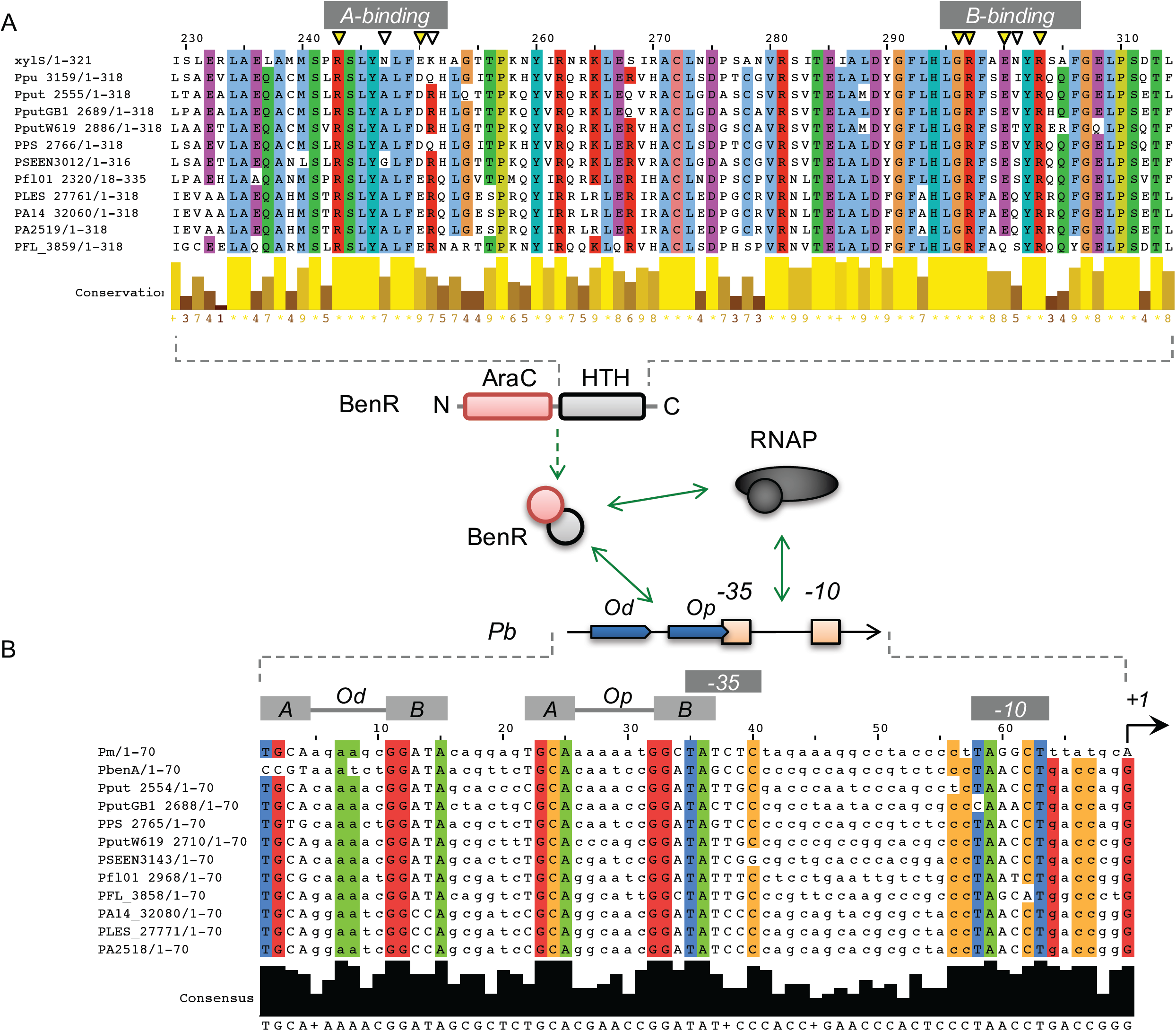
Analysis of BenR homologues and target promoters in *Pseudomonas*. **A)** Analysis of protein conservation between XylS from *Pseudomonas putida* mt-2, and BenR homologues from strains of *P. putida* (Ppu_3159, Pput_2555, PputGB1_2689, PputW619_2886, PPS_2765), *P. entomophila* (PSEEN3143), P. fluorescens (Pfl01_2968, PFL_3858) and *P. aeruginosa* (PA14_32080, PLES_27771, PA2518). Only the region relative to the HTH domain is shown. Critical aa for DNA recognition (labeled as A-binding and B-binding) are marked with inverted triangles, with conserved regions colored in yellow. On the middle, schematic representation of BenR interaction with the RNAP and the target promoter, shown the two binding sites (Od and Op) and the -35/-10 boxes at *Pb* and *Pm*. **B)** Promoter alignment for *Pm* from *P. putida* mt-2 and for *Pb* from several *Pseudomonas* strains. The two conserved boxes (A and B) from *Od* and *Op* binding sites are highlighted.

**Figure 2.**
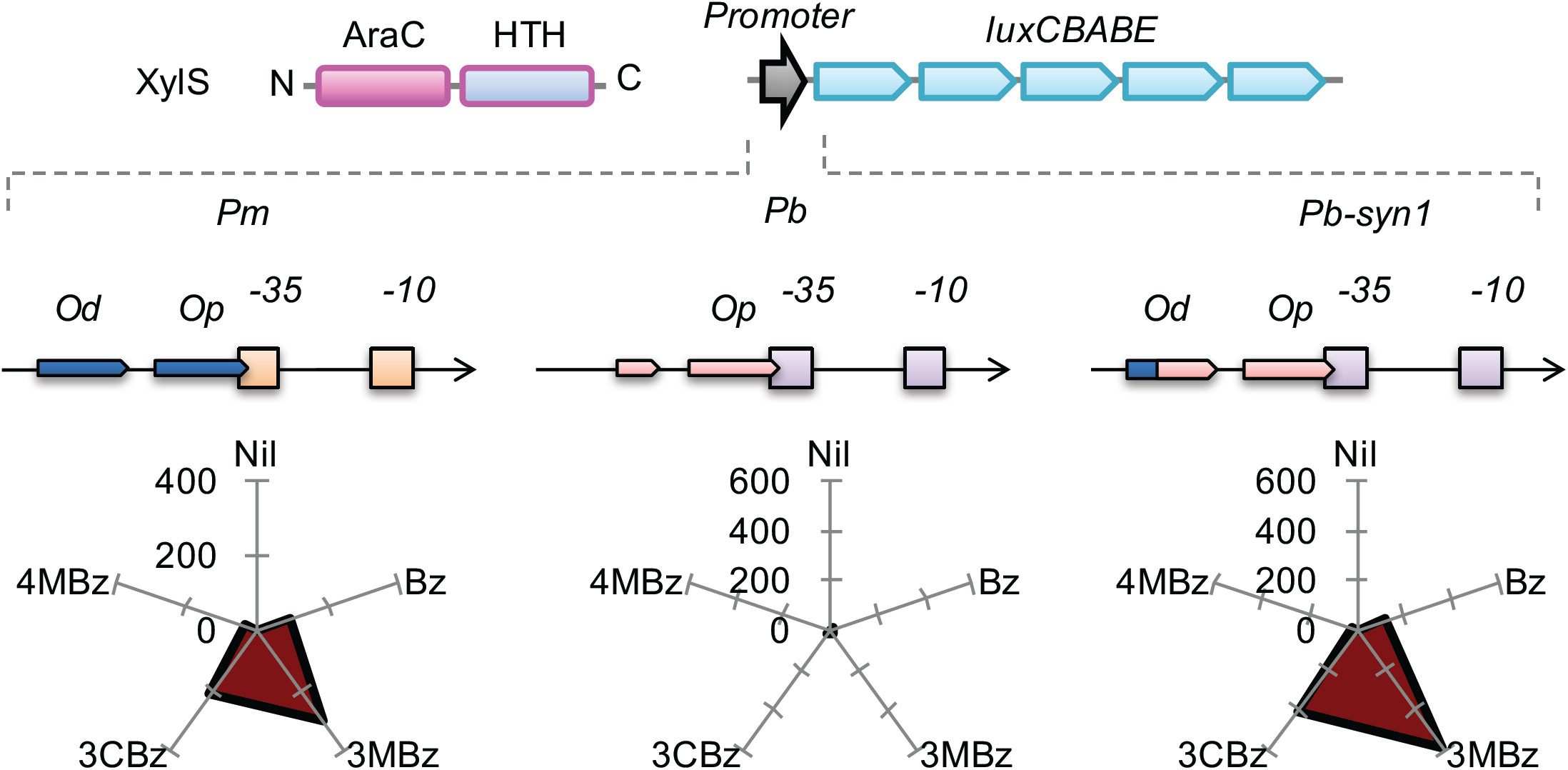
Recognition of *Pm* and *Pb* by XylS. For the analysis, *E. coli* DH5a strain was transformed with pSEVA438 (harboring a functional XylS expressed with its native promoter ^31^) and pSEVA226 (a reporter vector with the luxCDABE operon ^31^) harboring *Pm, Pb* or *Pb*-*syn1*, a variant of *Pb* endowed with the A box of Od from *Pm* ^25^. all strains were grown on minimal media to mid-exponential phase and then expose for 3 hours to 100 μM of benzoate (Bz), 3-methylbenzoate (3MBz), 3-chlorobenzoate (3CBz) or 4-methylbenzoate (4MBz). The results represent fold change of promoter activity relative to the non-induced condition.

### A single amino acid position is critical for aromatic recognition in BenR and XylS

After tracing critical differences in the DNA recognition requirements between BenR and XylS, we decided to investigate aa differences that could explain the divergence in the ligand selectivity between these two TFs. As presented before, BenR has a very narrow inducer selectivity as this TF can only respond to benzoate as inducer under natural conditions. However, XylS can be activated by a diverse collection of aromatic inducers, such as benzoate and methylated or chlorinated benzoate analogues ^27^. In order to gain insight into the molecular mechanisms responsible for these differences, we constructed a 3D protein structure for the N-terminal region of BenR using homology modelling. The resulting model was subject to molecular docking using benzoate, 3-methylbenzoate (3MBz), 4-methylbenzoate (4MBz) and salicylate (Sal). Using this approach, we obtained a protein structural model (**Fig. 3A**) and identified a potential cavity on the protein surface that accommodates well a benzoate molecule (**Fig. 3B**). Additionally, the potential binding pocket identified did not support well the binding of the benzoate analogues, suggesting a mode of inducer selectivity based on size exclusion. By analyzing the model and predicted binding pocket, we could identify 9 aa located from positions 67 to 121 that contributed to the surface of the cavity (**Fig. 3C**). With this information, we analyzed the conservation of these aa between BenR and XylS, hypothesizing that a change in some of these aa could explain the difference in ligand specificity between these two regulators. For our surprise, 8 out 9 aa from the predicted binding pocket were conserved between the two proteins. The only difference was at position 111, which is a Valine in BenR and an Alanine in XylS. Since Alanine has a shorter side chain, we hypothesized that this could lead to a wider binding pocket in XylS that could accommodate the methylated or chlorinated benzoate analogues. In order to investigate this possibility, we constructed point mutations in benR and xylS at aa positions 111 and 110, as we notice that this last position, while not involved in the binding pocket, was also not conserved between the two regulators (**Fig. 4A**). In addition to the point mutations, we constructed a chimaera between the N-terminal part of XylS and the C-terminal part of BenR and subjected this TF also to mutagenesis. All assays were performed using the cognate promoter for each TF (i.e., *Pb* for BenR and *Pm* for XylS) controlling a GFP reporter gene to allow investigation at the single cell level ^32, 33^ (**Fig. 4A**). As can be observed in **Fig. 4B**, wildtype BenR was specific to benzoate at all concentrations tested, while wildtype XylS displayed preferential response to 3MBz, an intermediated response to benzoate and lower response to 2MBz and 4MBz. When we mutated the position 110 of BenR from Pro to Ala or Gln (the aa found at this position in XylS), we could observe that the mutants presented the same expression profile of the wildtype but with reduced efficacy (**Fig. 4C**). However, when position 111 was changed from Val to Ala in BenR, the resulting protein variant did not display any response to the inducers tested. Contrary to the expected, changing Val at position 111 to Ala did not widen the inducer specificity of BenR. By the same token, the construction of BenR mutants with changes at both positions 110 and 111 resulted also in non-functional proteins (**Fig. S1**), potentially due to the role of Val111 on signal recognition. Taken together, these results indicate that Val111 is key for inducer recognition by BenR and strengthens the identification of the effector-binding pocket identified by homology modeling and molecular docking.

**Figure 3.**
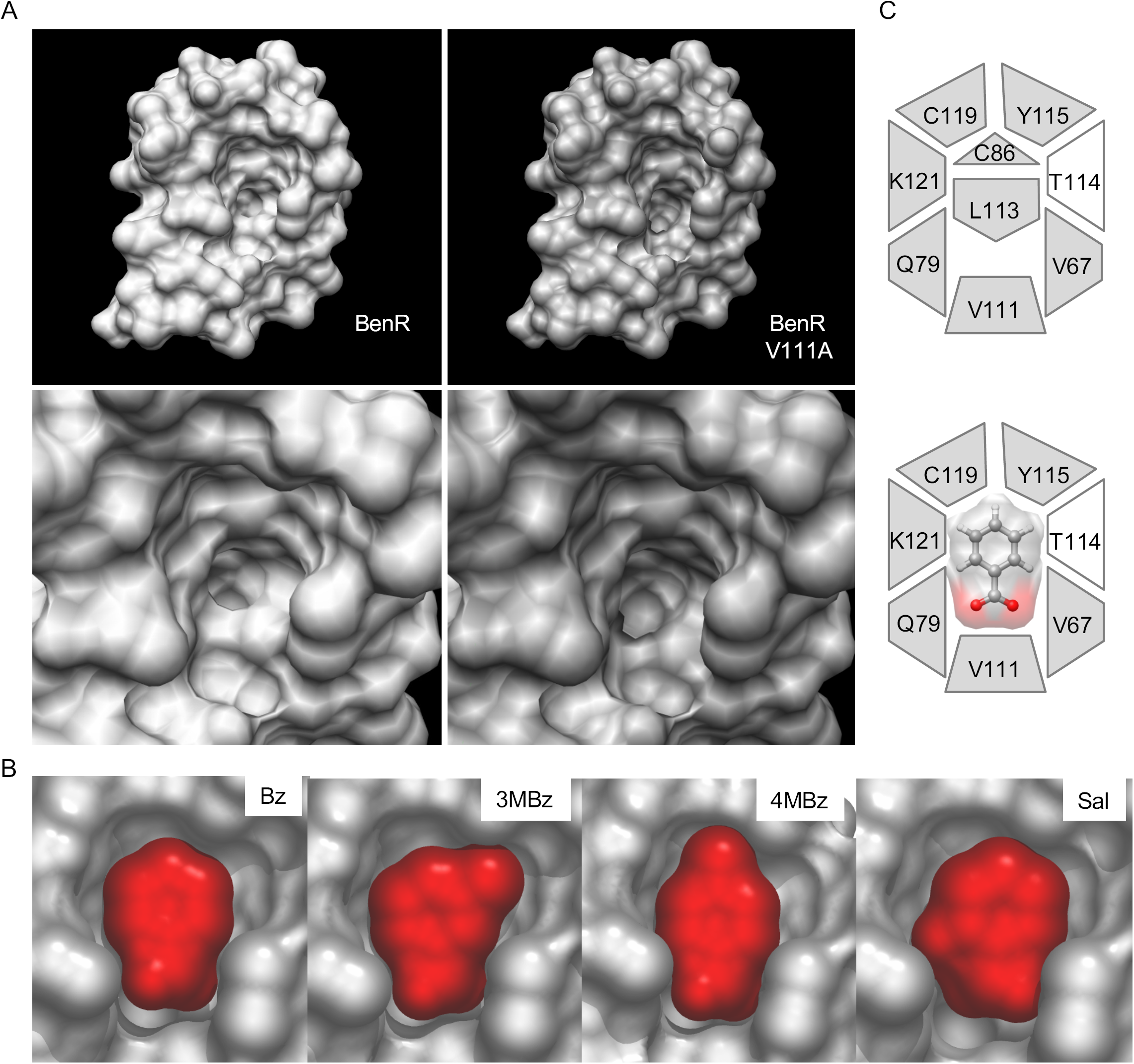
Mapping ligand binding site in BenR by protein structure prediction and molecular docking. **A)** Homology model of wild type (left) and V111A mutant (right) of BenR, with zoom at the larger cavity of both models shown on the bottom. **B)** Docking of benzoate (Bz), 3-methylbenzoate (3MBz), 4-methylbenzoate (4MBz) and salicylate (Sal) at the wild type BenR protein, with the molecules shown in red. **C)** mapping of the aa forming the potential binding site for benzoate at BenR (top) and the model for the interaction with the molecule in the bottom.

**Figure 4.**
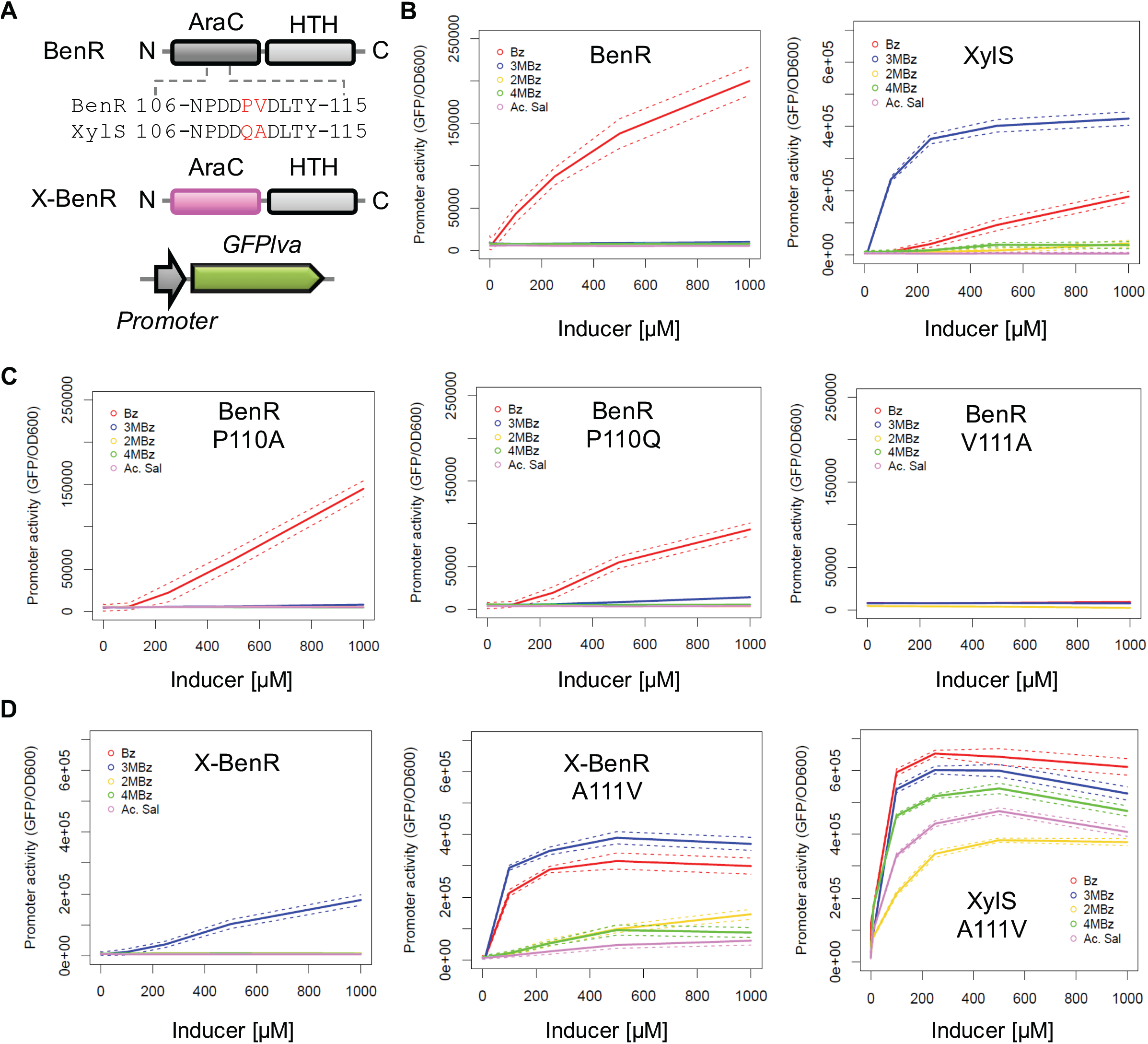
Experimental identification of critical aa for inducer recognition in BenR and XylS. **A)** Schematic representation of BenR and the chimeric protein X-BenR (a fusion between the N-terminal domain of XylS with the C-terminal domain of BenR), showing the non-conversed position 110 and 111 tested here. All protein variants were tested with a GFPlva reporter under the control of the cognate promoter (i.e., *Pb* for BenR and X-BenR and *Pm* for XylS). **B)** Analysis of promoter activity of BenR-*Pb* and XylS-*Pm* in response to different concentrations of benzoate (Bz), 3-methylbenzoate (3MBz), 2-methylbenzoate (2MBz), 4-methylbenzoate (4MBz) and acetyl salicylate acid (Ac. Sal). Solid line indicates the average from three independent experiments while dashed lines represents the lower and higher limits of the standard deviation. **C)** Analysis of promoter activity for BenR mutants at position 110 (BenR-P110A and BenR-P110Q) and 111 (BenR-V111A). **D)** Analysis of promoter acitivty for chimeric X-BenR protein and its variant with a mutation in postion 111 (X-BenR-A111V), as well as for the mutated version of XylS (XylS-A111V).

The construction of the chimeric protein harboring the N-terminal of XylS (the region responsible for ligand recognition) and the C-terminal of BenR (which recognizes DNA) resulted in the new protein X-BenR that displayed an induction profile similar to that of XylS but with a reduced efficacy (**Fig. 4D**). This result conformed that the required elements for inducer selectivity are placed in the first 196 aa of the proteins. In order to check the role of position 111 in the inducer selectivity of XylS, we constructed point mutants of XylS and X-BenR changing the Alanine at this position for a Valine. Unexpectedly, the resulting mutant proteins displayed an enhanced response to the optimal inducer (3MBz) as well as to the sub-optimal inducers benzoate, 2MBz, 4Mbz and salicylic acids (**Fig. 4D**). On the other hand, attempts to construct a chimeric protein harboring the N-terminal of BenR and the C-terminal of XylS (named B-XylS) resulted in non-functional products with no detectable response to benzoate or 3MBz (**Fig. S2**). Yet, single cell analysis of promoter induction by flow cytometry shows that while wildtype BenR and XylS-based expression systems presented clear unimodal patterns, the X-BenR chimaera and mutated version of XylS and X-BenR displayed a wider population distribution that could indicate stable sub-populations (**Fig. S3** to **Fig. S5**). Taken together, these results evidenced a remarkably different role of position 111 between BenR and XylS, which led to identification of two TFs variants with enhanced response to a wide range of benzoate derivatives.

### *A new set of TFs responsive to different aromatic compounds and Aspirin* ^*™*^

Building on the data above regarding inducer recognition of BenR and XylS, we developed novel TFs variants with reverse engineered ligand and promoter specificities. **Fig. 5** represents the overall performances of the new TFs in response to a set of benzoate derivate (**Fig. 5A**). As shown in **Fig. 5B**, BenR and X-BenR produced the lowest promoter output and are exclusively responsive to benzoate (BenR) or 3MBz (X-BenR). As these proteins have a BenR C-terminal domain, they can recognize both *Pm* and *Pb*. On the other hand, XylS has the intermediate level of promoter output and preference for 3MBz, followed by benzoate and only minor response to 2MBz and 4MBz. Yet, the mutated version of X-BenR (X-BenR-A111V) promoted an overall increased response to the suboptimal inducers of XylS and also response to Aspirin and ASA. Finally, the mutated version of XylS (XylS-A111V) display a tremendous gain of response to all benzoate derivatives tested, including Aspirin and ASA, with promoter outputs similar to that of wildtype XylS induced with their optimal inducer 3MBz (**Fig. 5B**). When fold-change is calculated relative to non-induced conditions, it can be noticed that maximal induction of XylS-A111V system (∼ 76 times) outcompetes that of wildtype XylS (∼ 69 times, **Fig. 5C**). In this new system, the response to ASA reach 42 and 45 times, respectively. Yet, as this enhanced response could be the result of the construction of a TF with a promiscuous effector specificity, we tested the response of XylS-A111V to toluene, xylene, phenol and a number of non-aromatic inducers (L-arabinose, fructose, glucose, IPTG and arsenite). As can be observed in **Fig. S6**, the system did not respond to any of these compounds, supporting the notion that only benzoate derivate could induce this mutated regulator.

**Figure 5.**
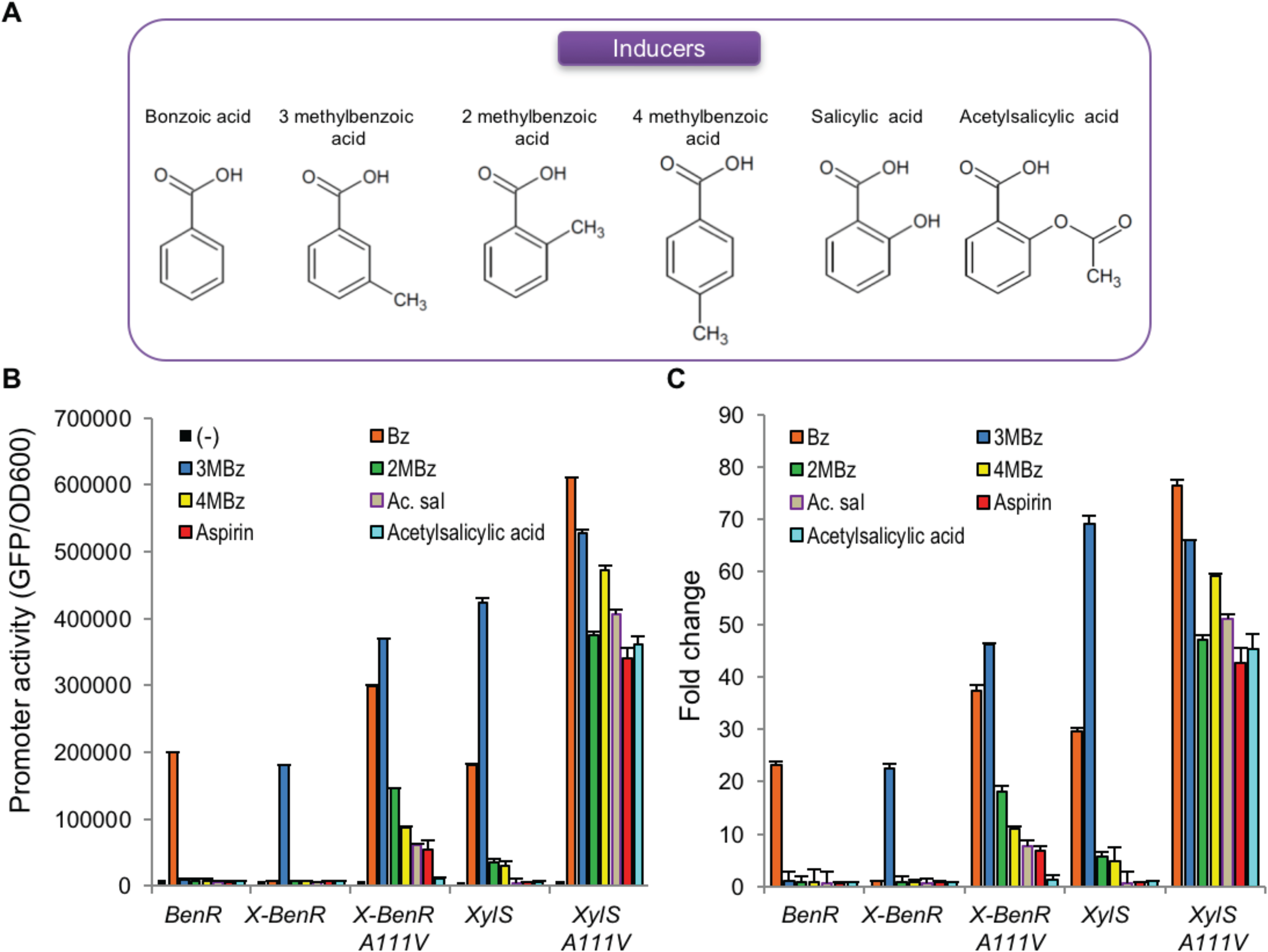
Analysis of transcriptional response to aromatic compounds and Aspirin. **A)** Schematic representation of the chemical structure of the different inducers tested. **B)** Promoter activity of the different proteins reported here after 3 hours of exposition of the different inducers. Aspirin refers to the commercial formulation obtained from a conversional drug store (with concentration adjusted to 100 μM), while acetylsalicylic acid refers to the pure compound obtained from Sigma-Aldrich. **C)** Fold-change calculated for each expression systemin response to the different inducers used. Error bars represent the standard deviation from thee independent experiments.

## Discussion

The results presented here shed some light on the molecular mechanisms for single recognition of BenR and XylS from P. putida. Of particular interest, XylS has been extensively investigated both on the context of the natural regulation of the meta pathways in *P. putida* mt-2 ^34, 35, 36^ as well as for its applicability as a universal expression system for gram-negative bacterium ^12, 37, 38^. Previous attempts have investigated the critical aa for inducer recognition and promoter activation ^12, 30, 39^, but these studies have not provided a clear molecular proposition on how this protein interacts with its ligands or the promoter. On the other hand, fewer studies have investigated BenR at the molecular detail ^24, 25, 26, 40^. As for the founds presented here, we initially expect that changes in the aa of the identified binding pocket of BenR could adjust the ligand selectivity of the protein, as reported for the TtgV regulator that can discriminate between molecules with one or two aromatic rings ^41, 42^. Yet, contrary to the initial size exclusion model proposed, our experimental validation of the binding site prediction supports a role for position 111 in both BenR and XylS as a key element connecting the binding of the ligand to the changes in domain arrangement of the protein, similarly to the mechanism of activation of AraC of *E. coli* ^43^. In this sense, the modelling of BenR with both N- and C-terminal domains suggest that the residue 111 is close to the interface between the ligand binding and DNA binding domains (**Fig. S7** and **S8**). In this scenario, Valine could make critical interactions in BenR that are disrupted when this aa is changed for Alanine. In the same way, changing Alanine in XylS (or in X-BenR) for Valine would allow the establishment of novel interactions that could enhance the performance of the protein, allowing the recognition of novel inducers such as salicylate or ASA. Previous work aiming the construction of XylS-*Pm* expression with increased response to 3MBz have reported several mutations in both the N- and C-terminal region of XylS. Among these mutants, an A111V mutation have been reported, but this mutation was selected together with one or two additional positions ^12^. Therefore, the role of position 111 in XylS has never been investigated in isolation.

It is interesting to notice that the mutations affecting signal specific of BenR and XylS either completely abolish protein activity or generated regulators with enhanced response to a series of ligand. in other words, it was not possible to switch the specific of the ligand-binding domains from one compound to another. This notion resembles the stem protein model investigated for XylR (another aromatic responsive regulator from *P. putida* mt-2), where selection of mutant proteins responsive to new ligands resulted in variants promiscuous to several aromatic compounds ^44^. It is also important to notice that the computational approach used here predict ligand binding pocket together with conservation analysis of phylogenetically related protein homologues represents a powerful tool to guide the rational design of TF variants. Similar approaches could be applied to other members of the AraC/XylS family of TFs, as well as regulators from different families, aiming the generation of novel expression systems for inducers of interest. While NahR, a LysR-type transcriptional regulator, is able to induce gene expression in response to salicylate ^10^, the TFs engineered here represent a new set of tools for the expression of genes of interest in response to salicylate and ASA. Additionally, the expression system based on XylS-A111V display a ∼ 10-fold change in response to 10 μM of ASA, in the range of the observed sensitivity for the natural ASA-responsive regulator NahR (which reaches a fold-change of 20 in response to similar concentration of the compound ^10^). These concentrations are in the range of the physiological concentrations of ASA in blood, as this molecules can reach levels as high as ∼30 μM after 20 min administration of the drug ^45^. Taken together, all results demonstrate the expression of the genetic toolbox for the engineering of synthetic circuits inducible to safe drugs.

## Material and methods

### Bacterial strains and growth conditions

The plasmids and bacterial strains used in this study are listed in **Table 1**. Cloning and assay procedures were performed in *E. coli* DH5α. All DNA manipulations, including cloning, PCR and transformations of *E. coli* were performed according Sambrook *et al*. ^46^. Bacterial strains were routinely grown in LB media supplemented with 36 µg mL-^1^ chloramphenicol or, when necessary, in M9 minimal media (6.4 g L-^1^ Na_2_HPO_4_•7H_2_O, 1.5 g L-^1^ KH_2_PO_4_, 0.25 g L-^1^ NaCl, 0.5 g L-^1^ NH_4_Cl, 2mM MgSO_4_, 0.1mM casamino acids, 1% glycerol) supplemented with chloramphenicol at 36 µg mL^-1^. Liquid cultures were shaken at 180 r*Pm* at 37°C during approximately 16h. The aromatic compounds used as inducers were all purchased from Sigma-Aldrich.

**Table 1.** Strains and plasmids used in this work.

| Strains and plasmids | Description | Reference |
| --- | --- | --- |
| **Strains** |  |  |
| *P. putida* KT2440 | *P. putida* mt-2 derivative | 49 |
| *E. coli* DH5α | F- φ80 ΔlacZ ΔM15 Δ(lacZYAA-argF) U169 recA1 endA1 hsdR17 R-M+ supE4 thi gyrA relA | 46 |
| **Plasmids** |  |  |
| pSEVA438 | Sm/Sp<sup>R</sup>, ori pBBR1; Expression vector harboring the *xyIS-Pm* expression system | 31 |
| pSEVA226 | Km<sup>R</sup>, ori RK2; reporter vector harboring the *luxCDABE* operon | 31 |
| pSEVA226-*Pb* | Km<sup>R</sup>, ori RK2; pSEVA226 with the *Pb* promoter cloned as a *EcoRI/BamHI* fragment | 25 |
| pSEVA226-*Pm* | Km<sup>R</sup>, ori RK2; pSEVA226 with the *Pm* promoter cloned as a *EcoRI/BamHI* fragment | 25 |
| pSEVA226-*Pbsyn1* | Km<sup>R</sup>, ori RK2; pSEVA226 with the *Pbsyn1* promoter cloned as a *EcoRI/BamHI* fragment | 25 |
| pMR1 | Cm<sup>R</sup>, ori p15a; dual mCherry/GFP<sub>lva</sub> promoter probe vector | 47 |
| pMR1-BenR-*Pb* | Cm<sup>R</sup>, ori p15a; pMR1 variant with *benR-Pb* expression system closed as a *BglIII/EcoRI* fragment | This study |
| pMR1-BenR-V111A | Cm<sup>R</sup>, ori p15a; pMR1-BenR-*Pb* with *benR* gene mutated at position V111A | This study |
| pMR1-BenR-P110A | Cm<sup>R</sup>, ori p15a; pMR1-BenR-*Pb* with *benR* gene mutated at position P110A | This study |
| pMR1-BenR-P110Q | Cm<sup>R</sup>, ori p15a; pMR1-BenR-*Pb* with *benR* gene mutated at position P110Q | This study |
| pMR1-BenR-A110A111 | Cm<sup>R</sup>, ori p15a; pMR1-BenR-*Pb* with *benR* gene mutated at positions P110A and V111A | This study |
| pMR1-BenR-Q110A111 | Cm<sup>R</sup>, ori p15a; pMR1-BenR-*Pb* with *benR* gene mutated at positions P110Q and V111A | This study |
| pMR1-X-BenR | Cm<sup>R</sup>, ori p15a; pMR1-BenR-*Pb* with *benR* with the N-terminal region replaced by that of *xyIS* | This study |
| pMR1-X-BenR-A111V | Cm<sup>R</sup>, ori p15a; pMR1-X-BenR with chimeric X-*benR* gene mutated at position A111V | This study |
| pMR1-XylS-*Pm* | Cm<sup>R</sup>, ori p15a; pMR1 variant with *xyIS-Pm* expression system closed as a *BglIII/EcoRI* fragment | This study |
| pMR1-XylS-*Pm*-A111V | Cm<sup>R</sup>, ori p15a; pMR1-XylS-*Pm* with *xyIS* gene mutated at position A111V | This study |
| pMR1-B-XylS | Cm<sup>R</sup>, ori p15a; pMR1-XylS-*Pm* with *xyIS* with the N-terminal region replaced by that of *benR* | This study |

### Plasmids constructions

The benR gene, *Pb*enR and *Pb* promoters were amplified by PCR using specific primers (**Table 2**) and *P. putida* KT2440 genomic DNA as template. The PCR products were digested with specific restriction enzymes (see underlined sequences in **Table 2**) and cloned into the *Pm*R1 vector ^47^, yielding the *Pm*R1-BenR-*Pb* (BenR) construction. BenR mutants: *Pm*R1-BenR-P110A (BenR-P110A), *Pm*R1-BenR-P110Q (BenR-P110Q), *Pm*R1-BenR-V111A (BenR-V111A), *Pm*R1-BenR-A110A111 (BenR-A110A111) and *Pm*R1-BenR-Q110A111 (BenR-Q110A111) were generated by CPEC site-directed mutagenesis methodology ^48^, using the *Pm*R1-BenR-*Pb* construction as template and the primers listed in **Table 2** (mutated base pairs are highlighted in bold and underlined). The xylS gene, Ps and *Pm* promoters were amplified by PCR using pSEVA438 vector as template ^31^, yielding the *Pm*R1-XylS-*Pm* (XylS) construction. XylS mutant *Pm*R1-XylS-A111V (XylS-A111V) was constructed by CPEC site-directed mutagenesis, using *Pm*R1-XylS-*Pm* construction as template and primers listed in **Table 2** (mutated base pairs are highlighted in bold and underlined). Two chimeric transcription factors were constructed. The first construction was generated by directly linking the N-terminal domain of XylS and the C-terminal domain of BenR using respectively *P. putida* mt-2 and KT2440 strains as templates. The second construction was generated by linking the N-terminal domain of BenR and the C-terminal domain of XylS using the *Pm*R1-BenR-*Pb* and the *Pm*R1-XylS-*Pm* as templates. All fragments were amplified by PCR. In the first construction, the *Pb*enR promoter was cloned upstream the chimaera and the *Pb* promoter was cloned upstream the GFPlva reporter gene yielding the *Pm*R1-X-BenR (X-BenR). The chimaera mutant *Pm*R1-X-BenR-A111V (X-BenR-A111V) was constructed using the CPEC and the *Pm*R1-X-BenR as template. In the second construction, the Ps and *Pm* promoters were cloned upstream the chimaera and the GFPlva reporter gene, respectively, generating the *Pm*R1-B-XylS (B-XylS). All PCR amplifications ware performed using Phusion High-Fidelity DNA polymerase (Thermo Fisher Scientific). All resulting constructions were sequenced using dideosyterminal methods in order to confirm the correct assembly prior to the fluorescence assays.

**Table 2:**
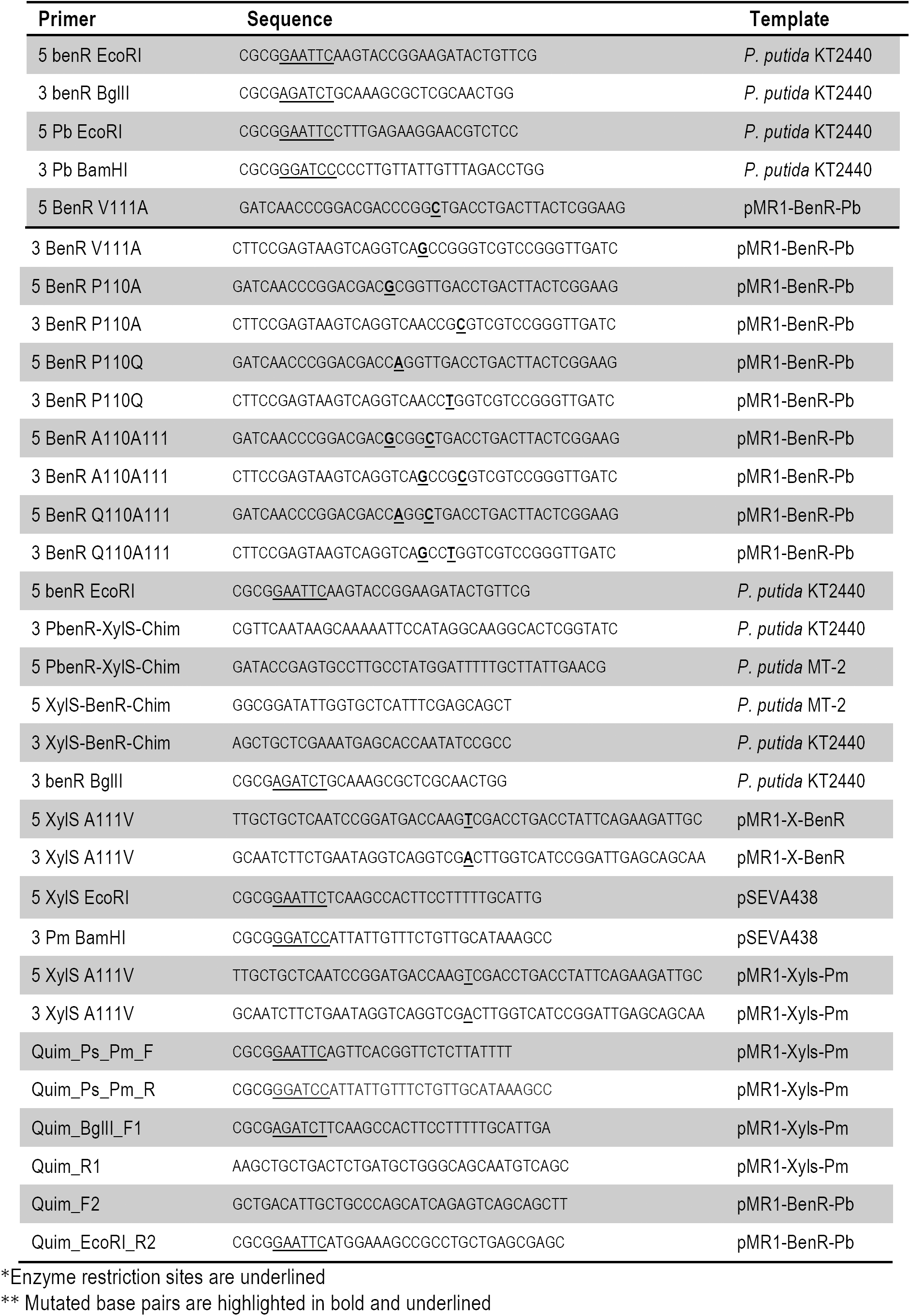
Primers used for each clone constructed in this work.

### GFP fluorescence assay and data processing

In order to measure the activity of all constructions performed in this work, plasmids were transformed into *E. coli* DH5α. Freshly-plated single colonies were grown in M9 minimal media supplemented with suitable antibiotics. The cultures (10 µL) were then assayed in a 96-well microplates with 170 µL of M9 minimal media and 20µL of the different compounds tested. When required, benzoic acid (Bz), 3-methylbenzoic acid (3MBz), 2-methylbenzoic acid (2MBz), 4-methylbenzoic acid (4MBz), salicylic acid (0-1000 µM); acetylsalicylic acid, Bayer^®^ Aspirin^*™*^ (1000 µM) and isopropil β-D-1-tiogalactopiranosida (IPTG), sodium arsenite, toluene, xylene, phenol, L-arabinose, fructose and glucose (100 µM) were used. Cell growth and GFP fluorescence were quantified using a Victor X3 plate reader (Perkin Elmer). The responsiveness of regulators was calculated as arbitrary units using the ratio between fluorescence levels and optical density at 600 nm (reported as GFP/OD_600_) or luminescence by optical density at 600 nm after background correction. As a control, all assays were performed without the addition of compounds as the threshold background signal during calculations. Fluorescence and absorbance measurements were taken at 30 min intervals up to 8 h. All the experiments were performed in technical and biological triplicates. Raw data were processed using ad hoc R script (https://www.r-project.org/).

### Flow Cytometry Analysis

High throughput single cell analysis of the bacteria carrying the BenR, XylS, XBenR, XylS mutant or XBenR mutant systems was run as follows. First, we selected single colonies of the transformed strain (*E. coli* DH5α) and grow overnight in M9 minimal medium (containing 6.4 g/L Na2HPO4-7H2O, 1.5 g/L KH2PO4, 0.25 g/L NaCl, and 0.5 g/L NH4Cl) supplemented with 2 mM MgSO4, 0.1 mM CaCl2, 0.1 mM casamino acids, chloramphenicol (36µg/mL) and 1% glycerol as the sole carbon source (supplemented M9), at 37°C and 180 r*Pm*. Next, overnight grown cells were diluted to 1:10 in fresh supplemented M9, and was grown for three hours, at 37°C and 180 r*Pm*. At this point, the cultures were induced with different concentrations (0µM, 10µM, 100µM, 250µM, 500µM and 1000µM) of the inducers. BenR system was induced with benzoate, XylS with benzoate and 3-metilbenzoate, X-BenR with 3-metilbenzoate, XylS mutant and X-BenR mutant with benzoate, 3-metil-Benzoate, Salicylic Acid and Acetyl-Salicylic Acid. After three hours of induction, the cultres were stored on ice and immediately analyzed for GFP fluorescence using the Millipore Guava EasyCyte Mini Flow Cytometer (Millipore). The results were analyzed by R scripts, using the flowCore and flowViz packages, available on Bioconductor (https://bioconductor.org/).

### 3D-structure model construction and docking analysis

The 3D models presented here were generated by SWISS-MODEL server (https://swissmodel.expasy.org/) using the best homologue for each protein. For the visualization of the models, PyMol (https://pymol.org/) and Chimaera (https://www.cgl.ucsf.edu/chimaera/) were used. In order to predict the potential binding site for the aromatic ligands, Swiss-Docking (https://www.swissdock.ch/) and Docking Server (https://www.dockingserver.com/web) were used. Additionally, Jalview (http://www.jalview.org/) was used for the visualization of protein and DNA alignments generated by T-coffee (http://tcoffee.crg.cat/apps/tcoffee/all.html).

## Acknowledgments

The authors are thanks to lab members for insightful discussion on this work. RS-R and MEG were supported by Young Research Awards, grant #2012/22921-8 and 2015/04309-1, São Paulo Research Foundation (FAPESP). LMOM, AS-M, LM-S and LFA were supported by FAPESP PhD Fellowships (grant #2016/19179-9, 2016/06323-4, 2017/17924-1 and 2018/04810-0).

